# Traveling waves shape neural population dynamics enabling predictions and internal model updating

**DOI:** 10.1101/2024.01.09.574848

**Authors:** S Mohanta, DM Cleveland, M Afrasiabi, AE Rhone, U Górska, M Cooper Borkenhagen, RD Sanders, M Boly, KV Nourski, YB Saalmann

## Abstract

The brain generates predictions based on statistical regularities in our environment. However, it is unclear how predictions are optimized through iterative interactions with the environment. Because traveling waves (TWs) propagate across the cortex shaping neural excitability, they can carry information to serve predictive processing. Using human intracranial recordings, we show that anterior-to-posterior alpha TWs correlated with prediction strength. Learning about priors altered neural state space trajectories, and how much it altered correlated with trial-by-trial prediction strength. Learning involved mismatches between predictions and sensory evidence triggering alpha-phase resets in lateral temporal cortex, accompanied by stronger alpha phase-high gamma amplitude coupling and high-gamma power. The mismatch initiated posterior-to-anterior alpha TWs and change in the subsequent trial’s state space trajectory, facilitating model updating. Our findings suggest a vital role of alpha TWs carrying both predictions to sensory cortex and mismatch signals to frontal cortex for trial-by-trial fine-tuning of predictive models.

---

The predictive coding framework proposes that the brain employs generative models to infer the state of the world based on prior sensory experiences^1–3^. These models are represented at higher-order levels of a cortical hierarchy, and predictions generated by the models are transmitted from higher-order to lower-order cortex through feedback connections. Any discrepancy between the predicted and observed sensory evidence leads to an error signal, which is transmitted along feedforward connections to update the models in higher-order cortex^3,4^.

There is substantial evidence of predictive cues influencing perception and behavior^5–7^, as well as neural oscillations influencing the signaling of predictions and errors^8–10^. However, the focus has been on stationary oscillations^9–11^, and not waves that propagate with flexible phase offsets across the cortex, known as traveling waves. Traveling waves may play a crucial role in cognitive processes^12–14^, including predictive perception^15^, by modulating neural excitability and the response to sensory input. Yet, the role of traveling waves in communicating predictions to lower-order cortex or in updating existing models after an error is unclear.

Another crucial question is how predictions emerge and develop in the brain during various stages of interaction with the environment^16,17^. Early in the learning process, the brain may rely on general priors and basic sensory information to generate predictions, resulting in a weak internal model. Over time, with increased exposure to the environment and error-driven model updating, the brain would be able to incorporate more nuanced contextual cues and refine its predictions, leading to the development of a strong internal model capable of making accurate predictions. Although computational modeling suggests trial-by-trial EEG event-related potentials (mismatch negativity) reflect prediction errors^18,19^, it is unclear how the neural representation of a weak internal model incorporates error signals and evolves into a strong model of the environment.

To investigate these questions, we used intracranial recordings in medically refractory epileptic patients performing an audio-visual delayed match-to-sample (A-V DMS) task, in which they learn the predictive value of sounds (which we validated previously in healthy volunteers)^10^. Using a dynamical systems approach, we found that learning about priors corresponds to changing trajectories in neural state space, where distance traveled correlated with trial-by-trial prediction strength. Furthermore, we show that stronger predictions are associated with anterior-to-posterior traveling waves in the alpha-frequency band. When the sensory evidence did not match predictions, lateral temporal cortex showed a phase reset of the alpha oscillation, greater alpha phase-high gamma amplitude coupling, and stronger high gamma power, a proxy for local neural firing^20^. This led to posterior-to-anterior propagation of alpha traveling waves to facilitate model updating, which manifested as a change in the subsequent trial’s state space trajectory. These findings shed light on the network-level mechanisms of how the brain learns and exploits statistical regularities.

## Results

### Predictions improved behavioral accuracy

We used an A-V DMS task to manipulate predictions and prediction errors (Fig. 1A). Participants first learned paired associations (A1-V1, A2-V2, A3-V3) between 3 sounds (A1, A2, A3) and 3 subsequent greeble images (V1, V2, V3) by trial-and-error. During learning of pairs, each sound and image had equal probability (33%) of appearing in any given trial, preventing participants from developing any differential predictions due to stimulus frequency. Following presentation of both stimuli, participants reported if the sound and image matched or not. After participants’ learned pairs, testing began. To manipulate participants’ predictions, we varied the probability of an image appearing after its associated sound. This probability was different for each sound: 85% chance of V1 after A1; 50% chance of V2 after A2; and 33% chance of V3 after A3 (Fig. 1A). Thus, A1 was highly predictive (HP), A2 was moderately predictive (MP), and A3 was not match predictive (NP). In our previous work^10^, we found that more predictive sounds led to faster reaction times and greater accuracy in match trials. In the current study, we could only measure accuracy because we included a second delay after image presentation in the task (withholding presenting response options until 1.6-1.8s after greeble onset), to allow characterization of the neural basis of prediction error processing. Nonetheless, we also found participants were more accurate when sounds had greater predictive value (F(2, 84)= 3.51, P= 0.0345) (Fig. 1B). These results demonstrate that the A-V DMS task manipulates participants’ predictions with corresponding changes in behavior.

**Figure 1.**
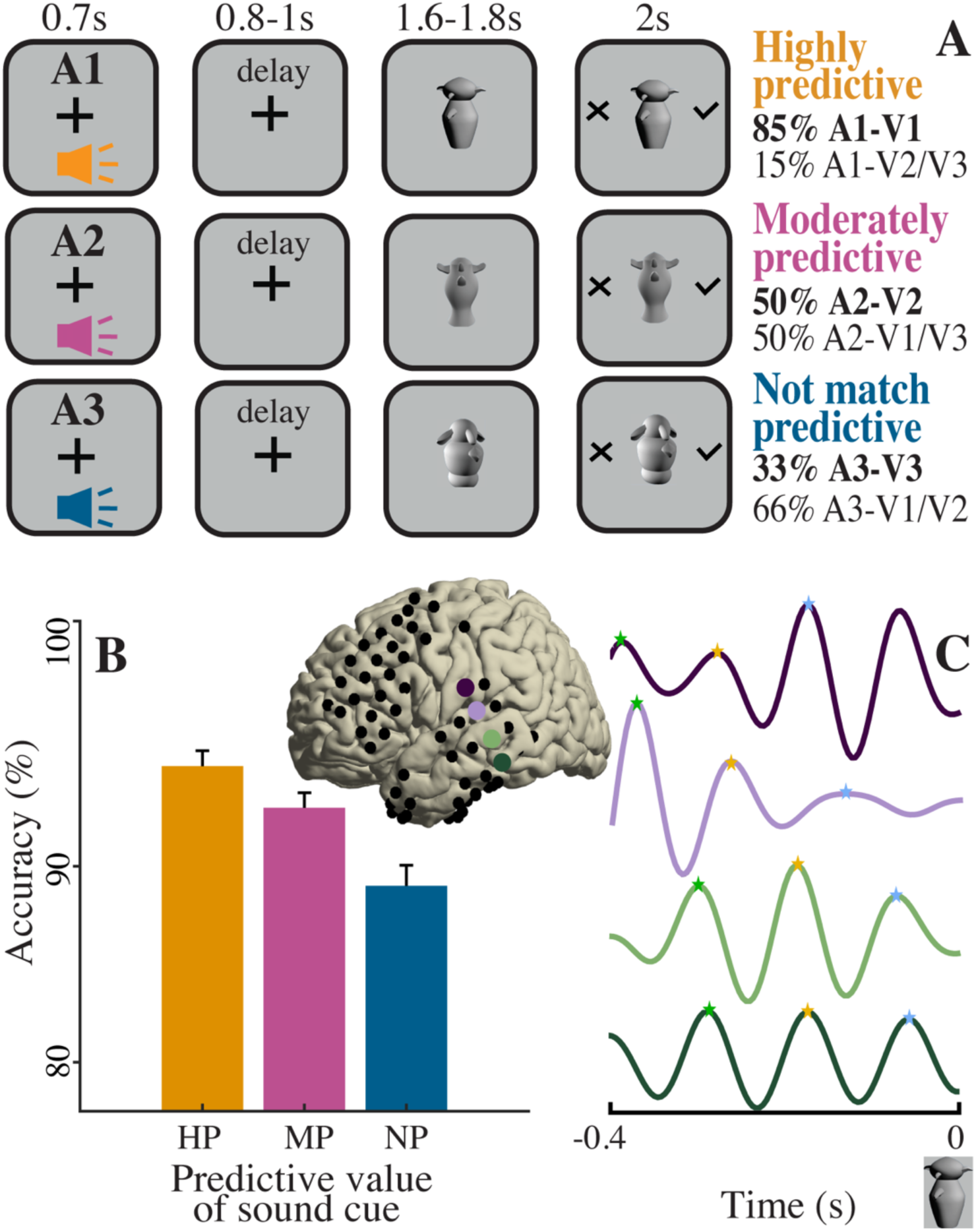
Experimental paradigm. (A) In the audiovisual delayed match-to-sample task, participants initially associated pairs of sounds and images in the learning phase, with all stimulus combinations equally likely. In the testing phase, we manipulated participants’ predictions by modifying the probability of an image appearing after its associated sound. (B) Population accuracy (%) of behavioral responses in highly, moderately and not-match predictive (HP, MP, NP, respectively) trials. Stronger predictive cues generated more accurate responses. (C) Alpha waves exhibiting propagation as a traveling wave across lateral cortex in an examplar participant. Four colored voltage traces in C correspond to colored electrodes on the brain inset. Top trace is anterior-most and bottom trace is posterior-most. Colored stars mark oscillation peaks, highlighting posterior electrodes lagging behind anterior ones.

### Predictions increased frontal high-gamma power and alpha-high gamma phase-amplitude coupling

We hypothesized that stronger predictions would have stronger representations in frontal cortex, evident as increased high gamma power (50-200Hz), which putatively reflects local spiking activity^20^. To test this hypothesis, we measured the activity of dorsolateral frontal cortical electrodes (8C, 8Av, i6-8, s6-8, SFL, 8BL, 9p, 9a, 8Ad, p9-46v, a9-46v, 46, and 9-46d according to Glasser et al.^21^ parcellation scheme) during the first delay period and discovered a positive relationship between the predictive value of the sound and high gamma power (effect was most pronounced for frequencies 60-100Hz; cluster-based permutation testing, P=0.03, Fig. S1A-C). Next, we investigated if this high gamma power change is modulated by the phase of low frequency oscillations. We found that phase-amplitude coupling in dorsolateral frontal electrodes was most pronounced at alpha frequency (around 9 Hz) modulating high gamma power (Fig. S1D-F; cluster-based permutation testing, P=0.0391). This suggests that predictive cues activate internal models and their predictions in frontal cortex.

### Anterior-to-posterior alpha traveling waves carry predictions

Alpha oscillations have been linked to predictive perception^9,10,22–25^, but the focus has been on stationary waves. We observed alpha oscillations traveling from the frontal to temporal cortex during the delay period after predictive sounds (Fig. 1C). To quantify the propagation of alpha oscillations (7-12Hz) across electrodes^26^, we computed the circular difference between the mean phase of the reference electrodes (in association auditory cortex, chosen due to all participants having coverage here) and that of each other electrode, at each time point. A positive or negative circular distance respectively indicates a leading or lagging electrode relative to the reference at each time point. To characterize the overall propagation of alpha waves across the entire delay period, we computed the circular mean of the circular distances across all time points within that period.

During the delay after HP sounds, there was a progression from relatively strong leading frontal electrodes through to lagging temporal electrodes (Fig. 2A) – compared to that from the delay after MP (Fig. 2B) and NP (Fig. 2C) sounds – suggesting that alpha oscillations start in the frontal cortex and propagate to the temporal cortex. To quantify the direction of propagation, we calculated the direction of the average spatial phase gradient across the electrodes at each time point; then averaged the directions across the delay period for the mean wave direction. This confirmed that the HP sound initiated directed alpha waves from frontal to temporal cortex (Rayleigh test of non-uniformity, Z=40.0540, P<0.0001; Fig. 2D, Fig. S1G and J). However, the propagation direction varied as a function of the predictive strength of the sound (Watson-Williams test, F(2, 300)=38.0469, P<0.0001; HP: Fig. 2D, Fig. S1G and J; MP: Fig. 2E, Fig. S1H and K; NP: Fig. 2F, Fig. S1I and L), and the distribution of propagation direction became more dispersed as predictive strength decreased (angular variance 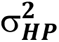 = 0.3703, 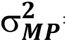 = 0.6076, 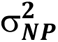 = 0.8292). Unlike the wave direction, the speed of alpha wave propagation did not vary according to the predictive strength of sounds (F(2, 234)= 0.001, P= 0.99; Fig. 2G-I). The median speed for each condition (range between 0.43 m/s to 0.57 m/s) was consistent with physiological constraints^27^ and previously reported values^13,28–30^.

**Figure 2.**
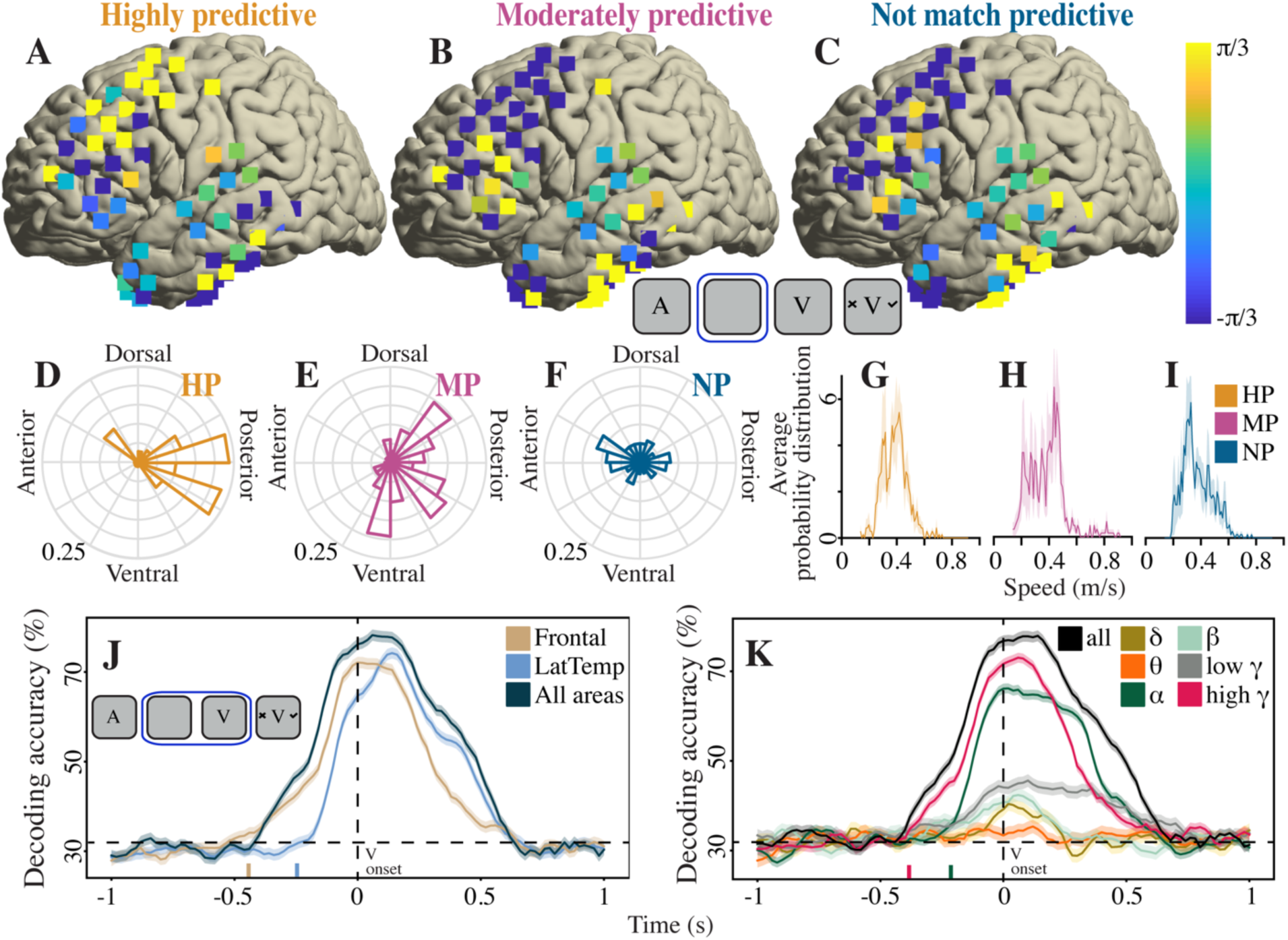
Anterior-to-posterior alpha traveling waves correlated to strength of predictive cue. A-C, In an exemplar participant, average circular distance of each electrode’s alpha phase from the spatial mean alpha phase, during delay period between sound and image, for (A) HP, (B) MP and (C) NP trials. D-F, Population probability distribution of alpha traveling wave directions across delay, for (D) HP, (E) MP and (F) NP trials. G-I, Population probability distribution of speed of alpha traveling wave: (G) HP, (H) MP and (I) NP trials. J and K, Decoder accuracy (%) classifying V1, V2 and V3 match trials when trained and tested at each time bin for (J) different brain areas (Frontal = dorsolateral, inferior, orbito-polar areas; LatTemp = lateral temporal areas; and All areas = all electrodes; area names based on Glasser et al.^21^ parcellation scheme) and (K) different frequency bands. Dashed vertical black line indicates greeble onset. Dashed horizontal black line indicates chance level decoding accuracy. Colored vertical bars on time axis in J and K indicate when decoding accuracy for corresponding condition first becomes significantly greater than chance. Insets in top row and J show task structure with blue outline highlighting time window of analysis for A-I and J-K respectively. A, sound cue; V, greeble image.

### Decodability of predictions emerges initially in frontal cortex, followed by lateral temporal cortex

If anterior-to-posterior alpha traveling waves carry predictions, then decoding accuracy should vary with electrode location, peaking first in frontal cortex where traveling waves initiate, and later in lateral temporal cortex that represents greebles^31–33^ (but prior to greeble onset). To test this, we used a time-generalized deep combinatorial recurrent neural network (RNN) to classify visual stimuli (V1, V2, V3) at successive time bins (0.02s) across match trials. Using either frontal (dorsolateral, inferior, orbito-polar) or lateral temporal activity from all frequencies (1-200 Hz) as input, the classifier was trained and tested across time from 1s before to 1s after greeble onset. The decoding accuracy of the classifier based on frontal activity became significantly greater than chance before the classifier based on lateral temporal activity (0.44s and 0.24s prior to greeble onset for frontal and lateral temporal cortex respectively; ANOVA; t=3.2; P=0.01; Fig. 2J). This follows the direction of the anterior-to-posterior traveling waves.

We then assessed the contribution of each frequency band from all electrodes to the decodability. Here we used Hilbert-transformed power estimates as input to the classifiers (time-generalized deep combinatorial RNNs, as before). The decoding accuracy for high gamma power, starting 0.38s before greeble onset, significantly exceeded chance, surpassing other frequency bands (Fig. 2K, red trace; ANOVA; t=4.5; P=0.002). Additionally, at 0.22s prior to greeble onset, the decoding accuracy for alpha power significantly exceeded chance and outperformed other frequency bands (ANOVA; t= 4.2; P=0.004), but to a significantly lesser extent than high gamma power (Fig. 2K, green trace; ANOVA; t=3.7; P=0.001). This suggests that high gamma and alpha activity reflect predictions.

### Prediction errors reset phase in lateral temporal cortex

In the predictive coding framework, a mismatch between a prediction and sensory input generates a prediction error signal, which plays an important role in updating existing generative models^1–3^. Because sensory inputs can induce phase reset or entrainment of cortical rhythms^34–36^, we tested whether prediction errors change the existing alpha oscillation phase in the lateral temporal cortex, where greebles are represented^31–33^. Since relatively few mismatches result from HP (A1) trials, for statistical robustness, we focused our analysis on the MP (A2) trials, which have an equal number of matches (no error) and non-matches (prediction error). Between 0.1s before greeble onset and 0.6s after, we assessed the consistency of phase estimates across trials for MP match (A2V2) and MP non-match (A2notV2, i.e., A2V1/A2V3) trials using inter-trial phase coherence (ITPC), a measure that ranges from 0 to 1. A value of 0 indicates no phase clustering, while a value of 1 indicates perfect clustering across trials. After greeble presentation, we found significantly stronger ITPC in the alpha band for MP non-match than for MP match trials. The effect was most pronounced within 0.2s after greeble onset (Fig. 3A and B, P<0.0001). Our ITPC results suggest that when there is a mismatch between prediction and sensory evidence, a phase reset occurs in sensory cortex.

**Figure 3.**
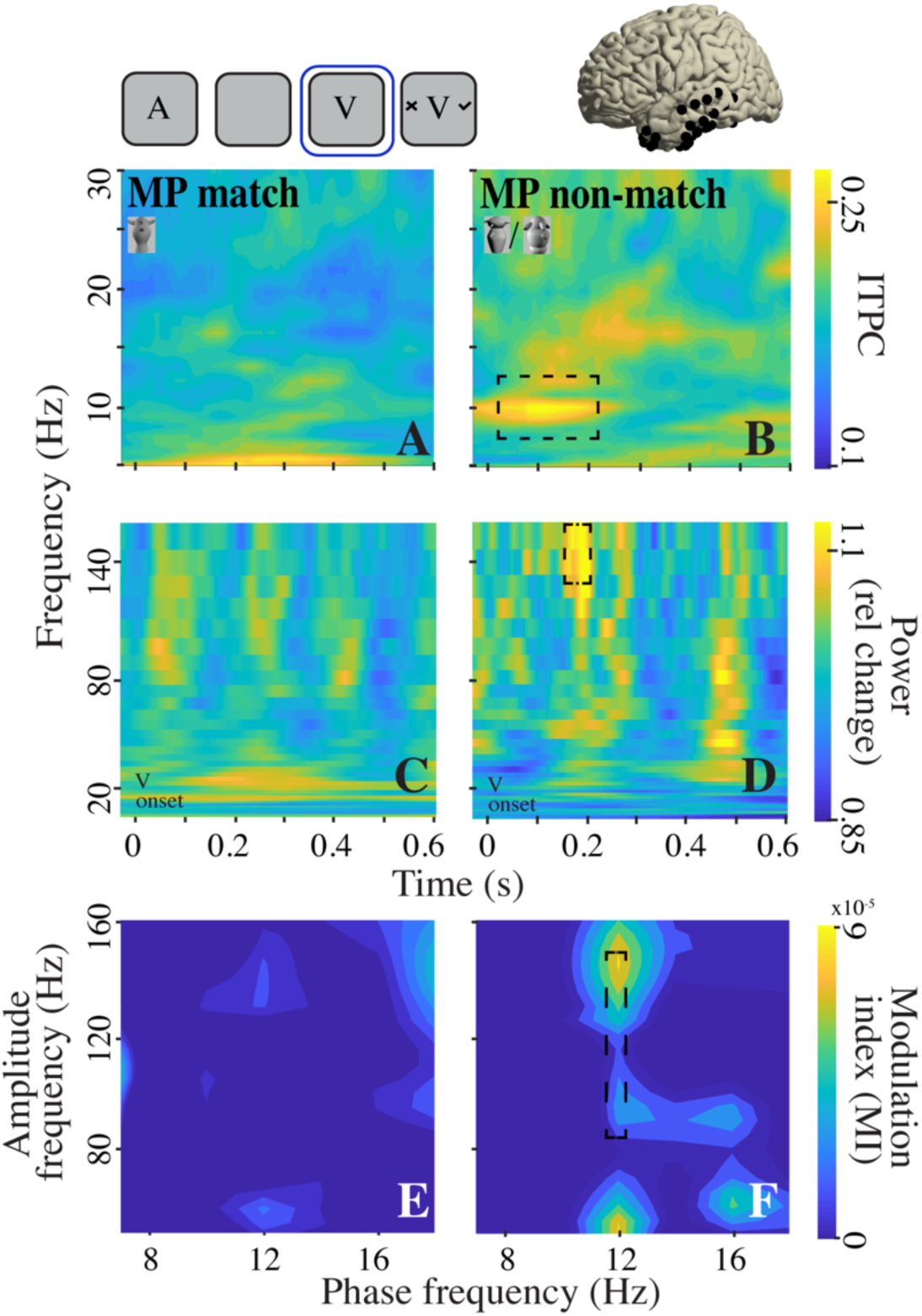
Prediction error increased ITPC, high gamma power, and phase-amplitude coupling in lateral temporal cortex. A-B, Population ITPC during period between greeble onset and response options for (A) MP match and (B) MP non-match trials. C-D, Population time-frequency power during period between greeble onset and response options for (C) MP match and (D) MP non-match trials. E-F, Population comodulograms for (E) MP match and (F) MP non-match trials. Most pronounced differences highlighted in dashed black squares after cluster-based permutation test for alpha-high gamma coupling. Plots in A-D aligned to greeble onset. Inset at top left shows task structure with blue square highlighting time window of analysis. A, sound cue; V, greeble image. Inset at top right shows lateral temporal electrodes in an example participant.

### Prediction errors increased high gamma power in lateral temporal cortex

Evidence suggests that prediction errors generate greater electrophysiological responses, measured as event-related potentials to unexpected stimuli in oddball paradigm in humans^37,38^ or spiking of neurons in macaques^9,39–43^. Based on these findings, one might expect error signaling to increase high gamma power, reflecting greater spiking activity in greeble-encoding neurons in lateral temporal cortex. We conducted a time-frequency decomposition comparing MP match and MP non-match trials and found a significantly larger increase in high gamma power after the greeble onset for MP non-match trials. The effect was most pronounced for latencies 0.15-0.19s and frequencies 131-160Hz (Fig. 3C and D; cluster-based permutation testing, P=0.028). These latencies were similar to those of single- and multi-unit neural responses to prediction errors^39–41^, consistent with increased high gamma power reflecting the generation of error signals.

### Alpha phase modulates high gamma power during prediction errors in lateral temporal cortex

We next investigated whether cross-frequency coupling plays a role in error processing, in light of the alpha phase reset and increased high gamma power. We calculated the modulation index (MI) for each possible pair of phase-modulating-frequencies (up to 30Hz) and amplitude-modulated-frequencies (50-200Hz) 0 to 0.6s after greeble onset. We found the phase of alpha oscillations strongly modulated high gamma power in MP non-match compared to MP match trials. Phase-amplitude coupling was most pronounced at 12Hz modulating 75Hz to 140Hz (Fig. 3E and F; cluster-based permutation testing, P=0.036). This result suggests that alpha phase reset and high gamma power changes are linked via phase-amplitude coupling. When there is a mismatch between prediction and sensory evidence, a phase reset occurs in the ongoing alpha oscillations in the lateral temporal cortex. This, in turn, triggers a phase-driven amplitude modulation of high gamma.

### Posterior-to-anterior alpha traveling waves carry error signals

To update internal models in the predictive coding framework, error signals are posited to propagate from posterior sensory areas to frontal cortex^1–3^. Given that traveling waves transiently modulate excitability as they pass^44,45^, and hence may influence synaptic plasticity, they could be useful for communicating error signals to frontal cortex for model updating. Therefore, we evaluated whether prediction errors initiate alpha traveling waves that propagate from posterior to anterior cortex (in line with proposed feedforward alpha waves^14,15^). We compared MP match trials (Fig. 4A), where predictions match sensory evidence, to MP non-match trials (Fig. 4B), where there is a mismatch and a need for model updating. Only in MP non-match trials, lateral temporal cortical electrodes led frontal electrodes after greeble onset (Fig. 4B), suggesting a posterior-to-anterior propagation of alpha waves. We calculated the direction of the average spatial phase gradient across the time window after greeble presentation (Fig. 2C and D) and confirmed that MP non-match trials trigger alpha waves from posterior-to-anterior cortex (Rayleigh test of non-uniformity, Z=4.19, P=0.01; Fig. 4D, Fig. S2B and D); whereas MP match trials still show anterior-to-posterior alpha waves (Rayleigh test of non-uniformity, Z=44.73, P<0.0001; Fig. 4C, Fig. S2A and C). The traveling wave directions significantly differed between MP match vs MP non-match trials (Watson-Williams test, F(1, 200)=100.18, P<0.0001). Our findings suggest that prediction errors trigger the propagation of posterior-to-anterior alpha waves, carrying error information to frontal cortex, potentially to update existing models.

**Figure 4.**
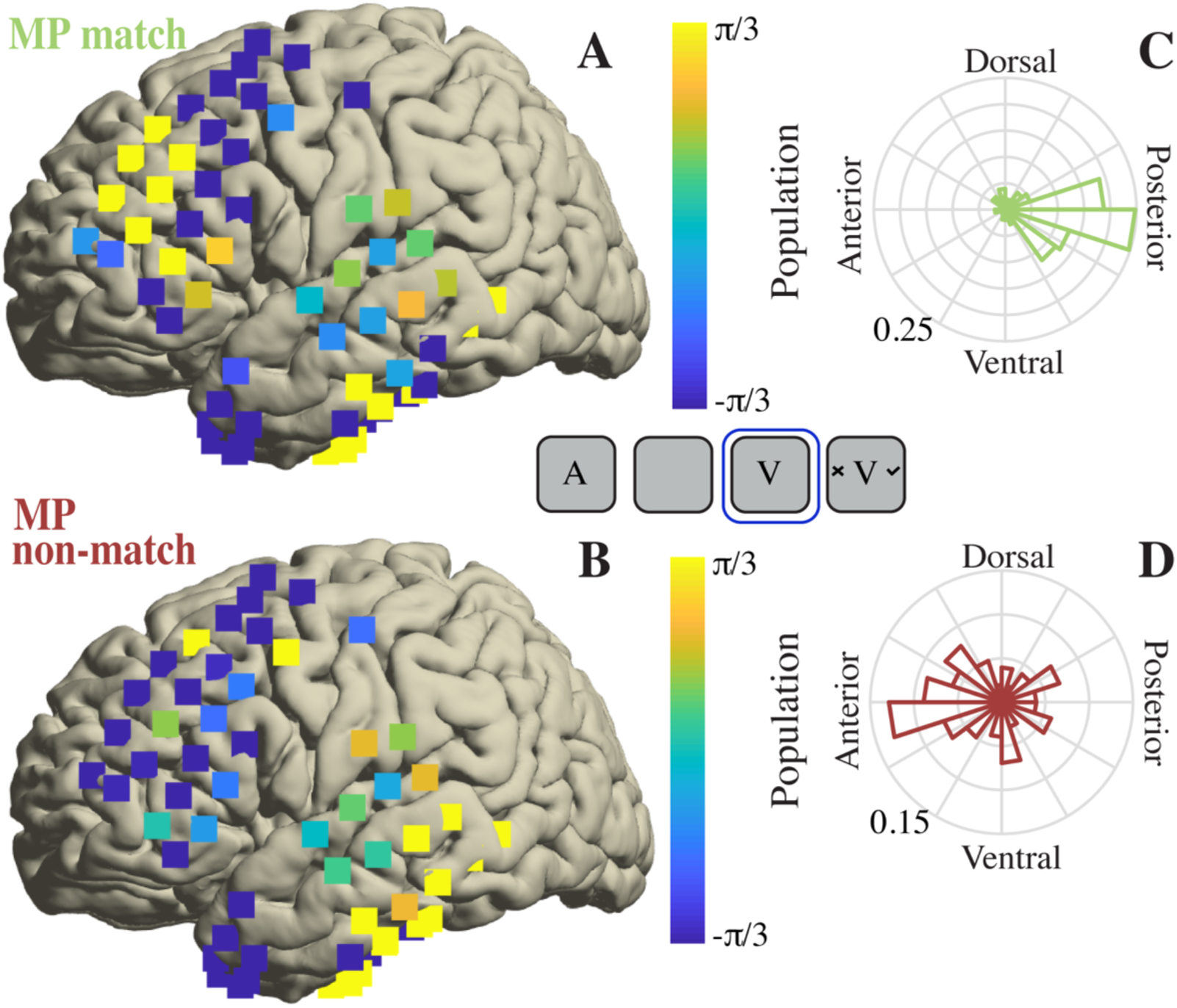
Posterior-to-anterior alpha traveling wave after prediction error. A-B, for an exemplar participant, average circular distance of each electrode’s alpha phase from the spatial mean alpha phase during period between greeble onset and response options for (A) MP match and (B) MP non-match trials. C-D, population probability distribution of alpha traveling wave directions across the same period for (C) MP match and (D) MP non-match trials. Inset shows task structure with blue square highlighting time window of analysis. A, sound cue; V, greeble image.

### Neural population state dynamics reflect prediction learning and model updating

Finally, we investigated how our brain acquires an accurate internal model of the world, to characterize how it progresses from an initially weak and inaccurate model to eventually generate precise predictions. All three sounds initially provided similar predictive information in our paradigm, but progressing through trials during testing, each sound developed a distinct predictive value: HP, MP or NP. We tracked how neural population dynamics evolved as sounds developed stronger predictive value. To do this, we used a dynamical systems approach. We divided our trials across a session into three equal phases: early, middle and late. In the early phase, the causal power^46^ – a measure of how likely a sound will be followed by its paired image, taking into account all stimulus relationships – of A1 (HP), A2 (MP) and A3 (NP) sounds overlapped (Fig. 5A). In the middle phase, the causal power started to separate but not completely. In the late phase, the causal power followed the order HP > MP > NP sounds. To capture the neural effects of prediction learning, we compared the early and late trial phases. We used the dimensionality reduction technique known as jPCA^47^, to project the high gamma power of all electrodes for each condition (HP, MP, NP) onto a lower-dimensional plane. We focused on high gamma power because it likely reflects local neural population activity changes more closely linked to prediction learning. We found that learning the predictive values of sounds, from early to late trials, corresponded to dynamic changes in trajectories within the jPCA space. Specifically, neural population activity for HP sounds (Fig. 5B) in late trials exhibited a distinct clustering pattern that was further away from early trials, compared with MP (Fig. 5C) and NP (Fig. 5D) sounds. Across all participants, the magnitude of this distance between early and late trials correlated with the predictive value of the sound (Fig. 5E; one-way ANOVA, F(2, 18)= 9.07, P=0.0019).

**Figure 5.**
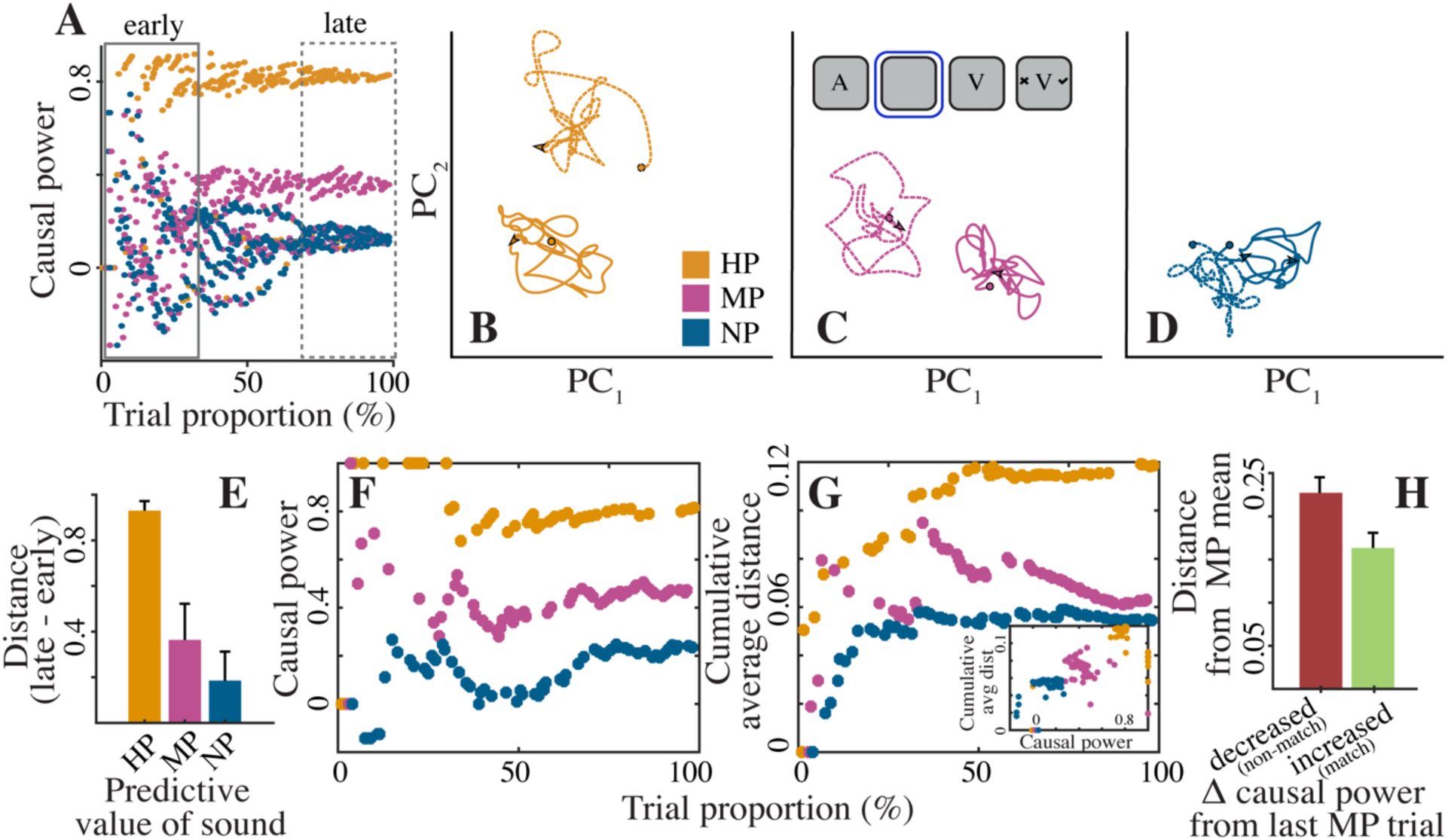
Distance traveled in state space correlated with the strength of predictive cue. (A) Population causal power as a function of the proportion of all trials. Solid box highlights early trials where causal power of HP, MP and NP overlapped; and dashed box highlights late trials where causal power followed the order HP > MP > NP trials. B-D, Mean neural trajectories for (B) HP, (C) MP, and (D) NP trials; solid lines, early trials, and dashed lines, late trials. Circle marks starting point, and arrow indicates final position (at greeble onset), of the trajectory. Inset in C shows task structure with blue square highlighting time window of analysis. A, sound cue; V, greeble image. (E) Population distance between early and late trajectories for HP, MP, and NP trials. (F) Causal power, and (G) cumulative average distance from the initial trial for each condition (HP, MP, NP) as a function of trial proportion, illustrated for an exemplar participant. Inset in G shows correlation plot illustrating the relationship between causal power and cumulative average distance for the exemplar participant shown in F-G. (H) Population distance of each MP trajectory from the mean MP trajectory influenced by changes in causal power from the preceding MP trial. Here, decreased (increased) causal power indicates a non-match (match) between prediction and sensory stimulus in preceding trial.

Because the distance traveled from the early trials depicts prediction learning, we hypothesized that the neural trajectory on a trial-by-trial basis depends on change in causal power, a metric that captures this learning. We calculated the distance of each trial from the first trial of the same condition. Our analysis revealed an alignment between the cumulative average distance and the fluctuations in causal power, reflecting a close association between these two variables. For example, in two participants shown in Fig. 5F-G and Fig. S3A-B, separation of HP (A1) and MP (A2) causal powers, and separation of the cumulative average distance, happened around the same trial proportion (40-50%). Furthermore, as shown in Fig. S3, this association between causal power (Fig. S3A) and cumulative average distance (Fig. S3B) also held in instances when the causal power of A3 (NP) was greater than A2 (MP) during the initial 0-50% trials (as was initially the case in this session due to pseudorandom stimulus presentation). To quantify these observations, we conducted a correlation analysis between causal power and cumulative average distance on a trial-by-trial basis (Fig. 5G inset, and Fig. S3C). Across all participants, we found a significant positive correlation between causal power and cumulative distance on a trial-by-trial basis (R(1159)=0.306, P<0.0001).

In the predictive processing framework, prediction errors play a crucial role in fine-tuning models. When sensory stimuli align with predictions, this leads to strengthening of the existing model; whereas mismatches prompt model adjustments to enhance predictions in subsequent iterations. To characterize the underlying neural mechanism, we analyzed how prediction errors in one trial impact the neural trajectory of the subsequent trial (for MP trials, which had equal numbers of match and non-match trials). We first established a neural representation of the model space for each participant, capturing the neural trajectory for the 0.1s prior to greeble onset in all MP trials (after a distinct separation of HP, MP and NP causal power), which we refer to as the ‘MP mean’. Using this as our baseline, we examined how individual MP neural trajectories deviated from this mean trajectory (calculation schematized in Fig. S3D) based on whether there was a prediction-sensory stimulus match (indicated by a causal power increase in Fig. 5H) or a non-match (indicated by a causal power decrease in Fig. 5H) in the preceding trial. We found that following a mismatch, the trajectory moved farther away from the mean trajectory, as shown by the larger distance from the MP mean in Fig. 5H (red bar). Conversely, following a match, the trajectory was closer to the mean trajectory, as shown by the relatively smaller distance in Fig. 5H (green bar). The neural trajectory distance from the “MP mean” was significantly different for preceding match versus non-match trials (one-way ANOVA, F(1, 404)= 7.72, P=0.0057). Taken together, our results capture how an initially imprecise neural representation of a model is iteratively fine-tuned by incorporating prediction errors (as well as alignments of the prediction and sensory stimulus). This suggests that predictive learning reflects an evolving neural dynamical system.

## Discussion

Our results show that acquiring knowledge about priors alters paths within the neural state space, i.e., paths shift towards a new subspace. The distance of this shift is correlated with the trial-by-trial prediction strength. Furthermore, we demonstrate that predictive cues activate frontal cortex, reflected in high gamma power, initiating alpha traveling waves that propagate from frontal to temporal cortex and carry predictive information. A mismatch between the prediction and the incoming stimulus triggers a reset in the alpha oscillation phase of the temporal cortex, leading to alpha phase-coupled modulation of high gamma power and the generation of a high gamma error signal within the temporal cortex. Subsequently, the alpha traveling wave direction reverses, now propagating from the temporal to frontal cortex to facilitate model updating, resulting in a change in the subsequent trial’s state space trajectory. Overall, our study suggests that traveling waves play a pivotal role in shaping neural population dynamics, to support predictive processing and model updating for a more precise representation of the world.

Previous modeling work^15^ has reported that priors can induce alpha traveling waves carrying predictive signals from anterior to posterior cortex, especially in the absence of sensory input like the delay period between sounds and greebles here. In our previous study^10^, using a paradigm similar to the current one, we observed that sound-induced predictions activate the corresponding greeble’s sensory representation before greeble onset. This is thought to occur through baseline activity increases in sensory neurons that are tuned to the expected stimulus^48–50^. Electrophysiological recordings from macaques have shown that traveling waves can modulate spiking activity, suggesting their capability to modify the baseline firing of neurons^45^. Taken together, traveling waves may modulate the baseline activity of greeble-encoding neurons in the lateral temporal cortex – reported location of greeble representations^31,32^ – to pre-activate sensory templates. Such pre-activations result in more efficient behavior^51,52^, as in our study, where participants performed more accurately for stronger predictive cues.

Modulation of oscillatory phase is a common phenomenon observed during attention to regulate responses to relevant stimuli^34,36^. Additionally, predictive cues have been reported to cause phase resets and biasing of phase to enhance stimulus processing^53,54^. Here we show that a mismatch between predictions and sensory evidence triggers a phase reset of alpha oscillations in lateral temporal cortex. Such phase resetting seemingly enables a break from feedback traveling waves and initiation of feedforward traveling waves upon registering an error signal. These findings suggest a link between attentional mechanisms and predictive perception. Attention may help recognize error signals when sensory evidence does not match predictions, which is key to updating internal models^1–3^. Specifically, error signals can be weighted through attention-like mechanisms, originating from, e.g., a subcortical source^4,55^, for model updating^56^ and improved predictive perception.

Feedforward communication is commonly viewed as operating through gamma frequencies^57,58^, including in the predictive coding framework^3^. This view is supported by Granger causality analysis of multi-electrode recordings from a cortical hierarchy^9^ showing increased gamma-band coherence along feedforward pathways for unpredictable stimuli. However, feedforward processing can also involve lower frequencies^59^. Alpha traveling waves propagate from posterior-to-anterior brain areas after visual stimulation^15^ and during memory encoding^14^. Because internally driven cognitive phenomena like memory recall can initiate anterior-to-posterior traveling waves in similar lower frequency bands^14^, the direction of the traveling wave may determine its functional role, not just the frequency *per se*. Consistent with this, we found anterior-to-posterior alpha waves during predictions, and posterior-to-anterior alpha waves during feedforward error signaling for updating existing models in higher-order cortex. Here, neural spiking activity representing a prediction or error signal would be modulated by the traveling wave along its direction of propagation.

There is growing evidence that predictive information is represented in a number of different areas across the brain, including the hippocampus, frontal lobe and sensory cortex^60–62^. These predictive representations may differ, e.g., distance in item sequence for the hippocampus versus spatial distance in visual cortex^60^. Further, the timing of the emergence of predictive models may differ, e.g., hippocampus early versus orbitofrontal cortex later^62^. Yet it remains unclear how predictive models and the latent structure of tasks are learnt. State space analysis techniques have been applied to understand artificial neural networks that mimic predictive learning in human behavior and decision-making^63^. Specifically, this modeling work has reported how learning predictive information changes the geometric properties of the latent space. Our trial-by-trial neural state space analysis provides empirical evidence for this, insofar that we show how iterative interaction with the environment can fine-tune a predictive model in fronto-temporal cortex, from initially weak and non-specific to eventually precise and accurate. Future studies need to further characterize the interrelationships between predictive representations in different cortical and subcortical areas, as well as their interdependence for predictive learning.

## Methods

### Human electrophysiology

Six neurosurgical patients who were diagnosed with medically refractory epilepsy participated in the study while they were undergoing chronic intracranial electroencephalography monitoring to identify potentially resectable seizure foci. Data were collected at University Hospital at University of Wisconsin – Madison and University of Iowa Hospitals and Clinics. Research protocols were reviewed and approved by the Institutional Review Boards at the University of Wisconsin – Madison and University of Iowa. Participants gave written consent. Research participation did not interfere with acquisition of clinically required data, and participants could rescind consent at any time without interrupting their clinical evaluation.

### Stimuli

We used biomorphic visual stimuli, known as greebles, from Michael Tarr’s laboratory (https://sites.google.com/andrew.cmu.edu/tarrlab/stimuli?authuser=0#h.i2ai1s51siku). Fig. 1A shows examples presented to participants. We used three grayscale greebles for each session, and each greeble was personified with a name. We used novel sounds (trisyllabic nonsense words) for the greeble names (e.g., “Tilado,” “Paluti,” and “Kagotu”)^64^. The sounds were generated using the Damayanti voice in the “text to speech” platform of an Apple MacBook. To avoid differences in the salience of stimuli, greeble images have similar size (13 degrees of visual angle (dva) in height and 8 dva in width), number of extensions and mean contrast, and greeble names have the same number of syllables and sound level (80 dB SPL).

### Audiovisual delayed match-to-sample task

Each trial involved the sequential presentation of a sound (trisyllabic nonsense word) followed by a greeble image. We refer to stimuli using the following notation: A1, A2 and A3 correspond to each of the three sounds used (A for auditory); and V1, V2 and V3 correspond to each of the greebles used (V for visual). Using this notation, audiovisual stimulus sequences containing the matching name and greeble are A1-V1, A2-V2 and A3-V3. Audiovisual stimulus sequences containing a nonmatching name and greeble are A1-V2, A1-V3, A2-V1, A2-V3, A3-V1 and A3-V2. We pseudo-randomized names for greebles (i.e., matching sounds and images) across participants.

#### Learning phase

During the first phase of the task, participants learn the association between the sounds and images (i.e., names of the greebles) through trial-and-error, by performing a match/nonmatch task. This phase is called the “learning” phase. Each trial starts with a blank blue screen (R=35, G=117, B=208, 1s duration; Fig. 1A). After that, a black fixation cross (size 1.16 dva; jittered 0.5s) is presented, followed by a sound, i.e., a greeble name voiced by the computer (0.7s duration). After a jittered delay period (0.8-1s), a greeble image is presented on the monitor screen for 1.6-1.8s. Following that, two symbols (✓ and X) were presented to the left and right of the greeble (9.3 dva from screen center). These symbols indicated participants’ two response options: match (√) or nonmatch (X). The symbol location, left or right of the greeble image, corresponded to the left or right response button, respectively: Z and M keys of a computer keyboard. Participants responded by pressing Z and M keys with their left and right index fingers respectively to submit a response. We randomly varied the symbols’ locations relative to the greeble image to minimize motor preparation (i.e., on some trials, a match response required a left button press and, on other trials, a match response required a right button press). In the learning phase, each greeble name and image had 33% probability of appearing in any given trial. This is to prevent participants from developing any differential predictions about the greebles because of greeble name or image frequency, during the learning phase.

#### Testing phase

Once participants show 80% accuracy in the learning phase of the task, they move on to the “testing phase” (200 trials). During the testing phase, we manipulated predictions by changing the probability of a greeble appearing after its learnt name. This probability is different for each greeble name and image. That is, in the testing phase, when a participant hears A1, there is 85% chance of V1 being shown (highly predictive, HP); when a participant hears A2, there is 50% chance of V2 being shown (moderately predictive, MP); and when a participant hears A3, there is a 33% chance of V3 being shown (not-match predictive, NP). This allows participants to make stronger predictions about the identity of the upcoming visual image after hearing A1, than after hearing A2 or A3, for instance. The task design involved creating the different predictive values for the three sounds in the testing phase, as well as balancing the stimulus numbers and match/non-match trials as closely as possible. That is, the testing phase of the task had a similar proportion of each voiced name and each greeble image, to control for stimulus familiarity, as well as a similar proportion of match and nonmatch trials to avoid response bias.

The behavioral task used in this study is the same as our previous work^10^, except that the response options were not simultaneously presented with the greebles. Here, the delay between presentation of greebles and response options allowed us to investigate processing of “error” signals in a predictive coding framework, distinct from action execution. To gauge any possible match bias, we previously ran “inversion trials” – requiring participants to report whether a greeble was upright or inverted – intermixed with match/non-match trials^10^. Since participants did not show a match bias, we only ran match/non-match trials to keep our current experiment run time under an hour. This allowed us to minimize any possible interference with clinical procedures, but still acquire a sufficient number of trials for our analyses.

### Behavioral data analysis

We analyzed only correct trials from the testing phase, reserving the learning phase solely to confirm participants’ acquisition of the correct sound-image associations. In order to focus on trials where the causal power of HP, MP and NP trials differed (Fig. 5C inset), we excluded the initial 50 trials from further analyses, unless otherwise mentioned. For the match trials (A1V1, A2V2 and A3V3), we calculated the overall accuracy for each trial type. To assess the predictive value of each sound cue on a trial-by-trial basis, we computed the causal power^46^: the amount of evidence that a sound “causes” a specific greeble, as opposed to a random different “cause.” We use “causal” in a statistical sense, capturing the relationship between the sound and closely following greeble. We updated causal power at each trial, to incorporate the extra information available^10^. Causal power values were the same for each stimulus at the start of the testing phase but, as participants performed more trials, causal power systematically differed between stimuli. This reflects the accruing information from the trial history.

### Anatomical reconstruction of electrodes

Participants underwent two imaging scans: a post-operative computed tomography (CT) scan, and a pre-operative or post-operative T1-MPRAGE structural magnetic resonance imaging (MRI) scan. We used FreeSurfer version 6.0 to segment the boundary between white and gray matter and extract cortical and subcortical labels^65–69^. We aligned CT and T1 scans using Freesurfer image analysis suite (v5.3) for participants at the University of Iowa, and using iElectrodes^70^ for participants at the University of Wisconsin – Madison. After scan alignment, we localized electrodes to the Human Connectome Project Multi-Modal Parcellation (HCP-MMP)^21^. Next, we mapped the parcellation into native space via fsaverage, to conform to the boundaries of the original FreeSurfer parcellations. We used the subcortical segmentation derived from FreeSurfer since the HCP-MMP does not label subcortical regions. We then extracted the distance of each electrode to the closest cortical label in the HCP-MMP (defined as the closest vertex on the boundary between white and gray matter) and to the closest subcortical voxel after excluding the ventricles and hypointensities. Depth electrodes were assigned an HCP-MMP or subcortical label based on the following criteria: i) If the electrode landed in an HCP-MMP or subcortical label, it was assigned to that label; ii) if the electrode was within 5mm of either an HCP-MMP label or subcortical label, and landed inside the brain, it was assigned to the closer label; or iii) if the electrode landed outside the brain, or was not within 5mm of a label, the electrode was excluded.

### Data processing

We performed all data processing using custom Matlab scripts and Fieldtrip software^71^. All recordings were first bandpass filtered 1 to 200 Hz. To remove 60 Hz electrical line noise, we then used a band stop filter of 59-61 Hz, 119-121 Hz and 179-181 Hz. Next, all recordings were down-sampled to 512 Hz, except for the travelling wave analysis, where recordings were down-sampled to 250 Hz. We excluded from data analyses any electrode channels in the epileptic region, showing epileptiform activity, or electrodes with excessive noise due to machine artifacts or poor contacts. For data from grid or strip electrodes, we re-referenced them to the common average. We used bipolar derivation to re-reference data from depth electrodes. Across all participants, 97 lateral temporal, 73 association auditory, and 106 frontal preprocessed electrodes were used in subsequent analyses.

### Traveling wave analysis

We used a 2-pass, 4th order Butterworth filter between 7-12 Hz to extract alpha band activity. We then applied a Hilbert transform (using ft_preprocessing.m function from Fieldtrip) to generate the analytic signal with instantaneous phase and amplitude of the alpha activity. Electrode coverage varied across our participants due to clinical considerations. To ensure we have sufficient and consistent coverage of electrodes in brain regions of interest across participants, we concentrated our analysis on electrodes on the lateral surface of cortex. To calculate spatial progression of alpha activity, we first calculated the reference phase at association auditory cortical electrodes at each time point: for grid electrodes, we calculated a circular average across all the association auditory electrodes; for depth electrodes, we chose the ventro-posterior most association auditory electrodes. We chose association auditory electrodes since all participants had more than two electrodes in association auditory cortices, thus giving similar reference electrodes across all participants. Then we computed the circular difference (using function circ_dist.m from CircStat Matlab toolbox^72^) between the mean phase and each electrode’s phase at each time point. Finally, we calculated the circular mean of this circular difference across all time points. If an electrode was leading an oscillation, it will have a positive circular difference in comparison to the reference phase; whereas if it was lagging, it will show a negative circular difference with respect to the reference phase. To quantify the direction of propagation of the alpha wave, we calculated the phase-gradient using Matlab’s gradient.m function on circular distances across a matrix. To apply the phase gradient function, each electrode’s data were put in a matrix representing the relative spatial location of the electrodes, such that the bottom right corner of the matrix had the ventro-posterior most electrode’s data. Additionally, for depth electrodes, the matrix size was constrained such that the number of columns corresponded to the number of probes in that participant. We calculated the average spatial phase gradient at each time point, for each participant, across all electrodes. This yielded the direction of alpha wave propagation at each time point. Our method was similar to that used by Halgren and colleagues^26^.

To calculate instantaneous propagation speed, we first estimated the instantaneous frequency by determining the first derivative of the instantaneous alpha phase at each time point. We then divided each channel’s instantaneous frequency by the magnitude of its phase gradient 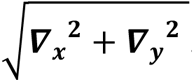 . This generated the instantaneous speed at each channel and time point. Then, for each time point, we calculated the median speed across all channels. We then generated a histogram of the speeds across time with bin width of 0.01 m/s for each patient and these histograms were then averaged across participants to generate distribution plots in Fig. 2G-I.

### Time-frequency decomposition

To investigate spectral properties during the first (the period between the sound and greeble image) and second delay (the period between the greeble onset and response options) periods, we time-frequency decomposed our “raw” neural data. We filtered data to multiple logarithmically spaced, partially overlapping passbands from 0.5–200 Hz^73^. Next, we applied the Hilbert transform to each filtered time series to compute the instantaneous amplitude of the analytic signal. The instantaneous power was then calculated by averaging across trials. All power measurements were baseline corrected, using the 0.4 s window prior to sound onset as the baseline.

### Decoding analysis

We used a combinatorial recurrent neural network (RNN) to classify V1, V2 and V3 trials relative to greeble onset, with Hilbert transformed power estimates as input. We ran seven different models, using the power spectral density of intracranial electroencephalography signals across six frequency bands, delta (1-3 Hz), theta (4-6 Hz), alpha (7-12Hz), beta (13-30 Hz), low gamma (31-49 Hz), and high gamma (50-200 Hz), as well as all frequencies (1-200 Hz) for the decoding analysis. For each model, we used a RNN with many-to-many architecture to decode visual stimuli using the power estimates from all our electrodes (pseudo-populated across all participants) in successive 0.02s time bins. The proposed RNN consists of three one dimensional convolutional layers (1-D CNN) of 32, 64, and 128 kernels with 7, 5, and 3 sizes respectively, followed by two Bidirectional long-short term memory (BiLSTM) layers of 64 and 128 neurons and 0.5 dropout, followed by a 8-neuron attention layer, a 128-neuron fully connected (FC) layer with 0.25 dropout, and a softmax layer as output. Considering x(t) as the input at time t, the output of the BiLSTM forward path is calculated as follows:

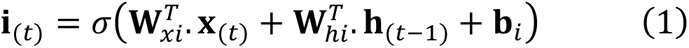

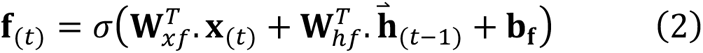

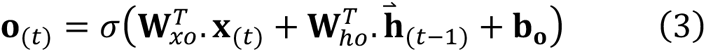

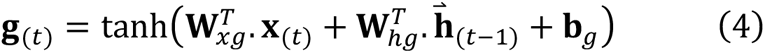

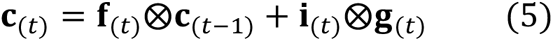

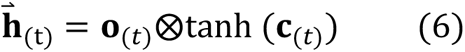

Where 𝐢_(.)_, 𝐟_(.)_, 𝐜_(.)_, 𝐨_(.)_ are the input gate, forget gate, cell gate and output gate respectively, 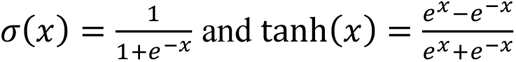 and the output of the backward path is 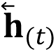. The attention layer output can be calculated as follows:

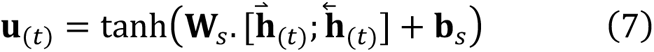

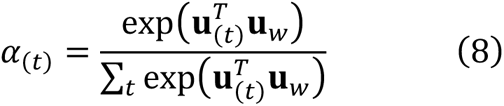

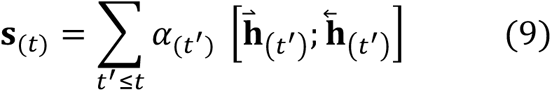

the output of the attention layer was fed to a fully connected layer with dropout followed by a softmax layer which results in class conditional probability as equation (10) specifies.

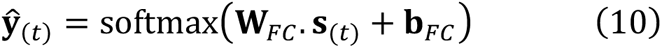

All CNN, BiLSTM, attention and FC layers’ weights and biases will be updated through backpropagation in time. Hyperparameters of the model including the number of hidden units for CNN, BiLSTM, Attention and FC layers, learning rate, dropout probability, learning rate of stochastic gradient descent, etc., are optimized via a grid search. The decoding accuracy is calculated with 10-fold cross validation with 60% of trials as the training set and 40% as the test set by shuffling and stratifying to avoid accuracy bias due to imbalanced classes.

To gain insight on the relative temporal dynamics of the above decoding performance, we used the same model as described in the previous paragraph. However, in this instance, we compared power estimates across all frequencies in different brain areas. Specifically, we used either frontal (dorso-lateral, inferior, and orbito-frontal) or lateral temporal areas, as inputs to the models.

In evaluating the efficacy of our model, decoding accuracy was ascertained using a 10-fold cross-validation method. Here, 60% of trials were used as the training set and the remaining 40% as the test set. The stratification and shuffling techniques were employed during this division of data sets, ensuring that we minimized any potential accuracy bias that could arise due to imbalanced classes.

To ensure a thorough and unbiased selection of trials, pseudo-population data generation was conducted 10,000 times. Accuracy measurements for test data were performed for each cross-validation fold and for each pseudorandom instance of the dataset, providing us with a robust statistical overview of the model’s performance.

### Inter-trial coherence (ITPC)

We divided the complex signal of our time-frequency decomposed data by the amplitude of the signal. Next, we summed the resultant angles across trials to calculate inter-trial phase coherence. In order to compute an unbiased estimate, we adjusted for the number of trials in each condition by dividing it by the number of trials (code adapted from https://www.fieldtriptoolbox.org/faq/itc/). We also calculated Raleigh-corrected ITPC scores, a popular ITPC measure to normalize for the number of trials (code adapted from https://mikexcohen.com/lecturelets/itpc/itpc.html). We found similar results using both measures.

### Phase-amplitude coupling (PAC)

We decomposed zero-meaned raw time series to complex analytic signals using the Hilbert transform as described above. We measured cross-frequency PAC using the modulation index (MI)^74^. We analyzed data between 0 to 0.6s after stimulus onset (aligned to greeble onset for Fig. 3E-F and aligned to sound onset Fig. S1D-F), using open source code from Adriano Tort’s lab (https://github.com/tortlab/phase-amplitude-coupling). We refer to the lower frequency oscillation, whose phase modulates the amplitude of the higher frequency activity, as the “phase-modulating frequency”; and we refer to the higher frequency activity as the “amplitude-modulated frequency”. We calculated the MI for each pair of frequencies, by binning the phase of the phase-modulating frequency into eighteen 20° intervals ranging from 0 to 360°. Subsequently, we determined the mean amplitude of the amplitude-modulated frequency within each phase bin. To transform the mean amplitude per phase into a probability distribution-like function, we normalized the resultant amplitude by dividing the mean amplitude in each phase bin by the sum across all bins. To quantify the deviation of the phase-amplitude distribution from the uniform distribution, we computed the Kullback-Leibler (KL) distance^75^. Finally, we calculated the MI by dividing the KL distance by the logarithm of the number of phase bins, which ensured the normalization of the measure to a range of 0 to 1. A MI value of 0 indicates uniform mean amplitude across all phase bins, signifying the absence of phase-amplitude coupling. As the amplitude distribution deviates further from the uniform distribution (as indicated by the KL distance), the MI value increases. We generated comodulogram plots (Fig. 3E-F, Fig. S1D-F) to illustrate the MI for different PAC frequency pairs: the phase-modulating frequencies correspond to 4-Hz bandwidths, incremented in 2-Hz steps; and the amplitude-modulated frequencies correspond to 10-Hz bandwidths, incremented in 5-Hz steps. To infer that the observed MI value was above chance level, we applied the “split-invert-splice” method used by Scheffer-Teixeira and Tort^76^. We kept the phase time series intact, while randomly splitting the amplitude time series. Then we concatenated the two parts inverted and calculated the comodulogram for the inverted time series. Actual MI values were corrected – actual minus inverted – to generate Fig. 3E-F and Fig. S1D-F.

### Causal power

We calculated causal power^46^ using the methods and code reported in our previous publication^10^. In our paradigm, causal power captures the amount of evidence that a sound “causes” a particular image, as opposed to a random different “cause.” Here, “causal” is used in a statistical sense, capturing the relationship between the initial sound and the image that closely follows in time.

### jPCA

We measured neural population-level dynamics using jPCA^47^. This involved projecting baseline-corrected high gamma power of all our electrodes for individual participants, during the delay period between the sound and image, to a lower dimensional space. We employed the process of "soft" normalization (parameter value = 10), where electrodes exhibiting strong responses were brought down to a range close to unity, while electrodes with weaker responses were limited to a range below unity. Further, we subtracted the average response across conditions from the response for each specific condition. We set the number of principal components to 6 for our analysis. To analyze the dynamic nature of prediction learning, we compared early versus late trials. First, we chose the jPCA plane that explained maximum variance for each patient. Subsequently, in that plane, to compare destinations of neural trajectories at the end of the delay, for each condition (HP, MP and NP), we computed the distance between the Cartesian coordinate of the "early" cluster and the "late" cluster during the time window 0.1s to 0s prior to greeble onset. The average distance for all participants is shown in Fig. 5E.

To investigate trial-by-trial changes in neural trajectories (Fig. 5G, and Fig. S3B), for each condition, we calculated the distance of each trial from the first trial of the same condition using the distance measure described in the above paragraph. Subsequently, to delineate dynamic changes in the distance measure relative to trial proportion, we transformed the distance metric into cumulative average distance, with the trial proportion dictating the chronological order.

To capture how a prediction error in a trial affects the neural trajectory of the subsequent trial, we first established a neural representation of the model space for each participant. To define this, we calculated the mean 0.1s (50 samples) neural trajectory prior to greeble onset of all MP trials presented after a distinct separation of A1 (HP), A2 (MP) and A3 (NP) causal power (i.e., 100^th^ trial onwards in a 200-trial block; the last half of the block of trials was chosen to ensure that there was no overlap in HP, MP and NP causal power). We call this mean trajectory “MP mean”. Next, for each MP trial’s 0.1s neural trajectory prior to greeble onset, we computed the point-by-point distance from the “MP mean” trajectory (as schematized in Fig. S3D inset) and averaged them, to generate a distance measure for each participant. Further, we calculated the difference between the causal power for a given MP trial and the preceding MP trial. A mismatch in the preceding trial resulted in a decrease in causal power; whereas a match led to an increase in causal power.

### Statistical analysis

We performed an ANOVA to compare the accuracies of match trials. In this analysis, the predictive value of the sound cue (HP, MP or NP) served as the independent variable, while accuracy was the dependent variable.

To assess the distributions of directions for travelling waves over time for HP, MP and NP trials during delay one (the period between the sound and greeble image) as well as for MP-match and MP-non-match trials during delay two (the period between the greeble onset and response options), we employed a Watson-Williams test (circ_wwtest.m). This test is akin to conducting a one-way ANOVA for circular data. To assess non-uniformity of traveling waves’ directions. we used a Rayleigh test (circ_rtest.m).

We employed a cluster-based permutation technique^77^ to compare conditions for ITPC, time-frequency power and PAC findings. To investigate ITPC differences between MP-match and MP-non-match conditions, we tested for differences in alpha frequency within the latency range of 0 to 0.6s after the onset of greeble stimuli. To compare the time-frequency spectra for the HP, MP and NP conditions during delay one in dorsolateral frontal cortical electrodes, as well as for the MP-match and MP-non-match conditions during delay two in lateral temporal cortical electrodes, we focused our analysis on the 80-200 Hz frequency range. For investigating prediction-related time-frequency changes, we concentrated our analysis on delay one (-0.8 to 0s before greeble onset); while for error-related activity, we examined the period from 0 to 0.2s following the onset of greeble stimuli. Finally, to compare PAC between MP-match and MP-non-match conditions, we selected the alpha band as our phase-modulating frequency of interest and the high-gamma band as our amplitude-modulated frequency of interest. To compare the distance between early and late neural trajectories across HP, MP, and NP conditions (Fig. 5E), we employed a one-way ANOVA test (anova1.m function in MATLAB). We also used a one-way ANOVA test to explore the impact of changes in causal power from the preceding MP trial on the subsequent MP trial’s trajectory (Fig. 5H).

To assess the decoding accuracy across different frequency bands and regions of interest (frontal vs lateral temporal vs all areas), we conducted a comparison involving 100,000 instances of accuracy. This analysis incorporated a ten-fold cross-validation approach with 10,000 pseudo-population randomization iterations using ANOVA across time. To account for multiple comparisons, we applied Holm’s correction.

Subsequently, we calculated the latency for each frequency band and regions of interest. This calculation relied on the decoding accuracy results to identify the time point at which each frequency band or regions of interest exhibited decodable information, signifying a significant departure from chance levels. These latency values were then compared across time using ANOVA with Holm’s correction to address multiple comparisons.

## Acknowledgements

This work was supported by: R01NS117901 grant to Y.B.S. and R.D.S.; Wisconsin Alumni Research Foundation Fall competition grant to Y.B.S.; Schwartz Fellowship, Department of Psychology, University of Wisconsin-Madison to S.M.; K23NS112473 grant to M.B.; R01DC004290 and R01GM109086 to K.V.N.. We thank Joel Berger, Haiming Chen, Christopher Garcia, Matthew A Howard III, Hiroto Kawasaki and Beau Snoad for expert assistance.

---

**Figure S1.**
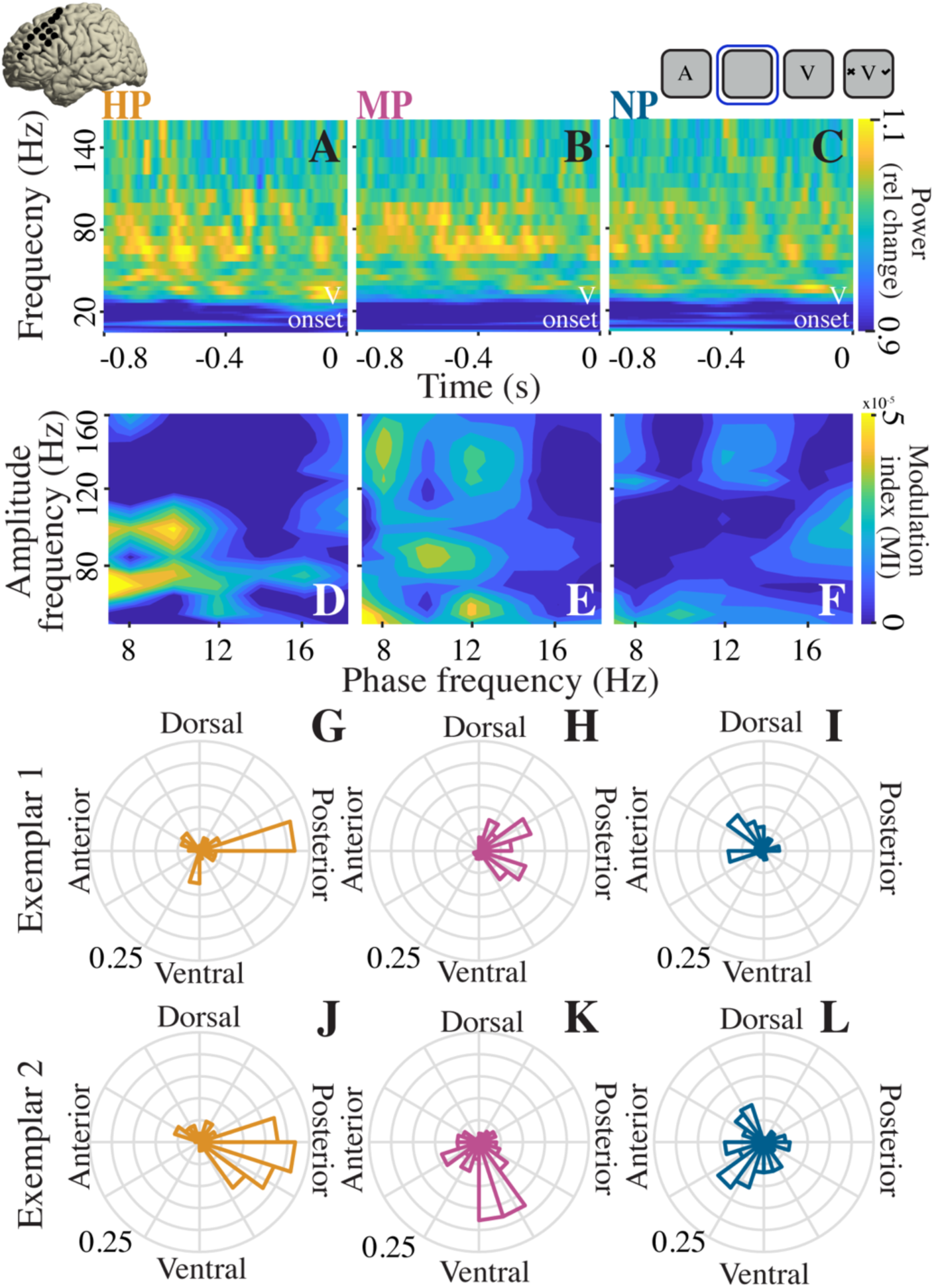
High gamma power and alpha-gamma phase-amplitude coupling in dorsolateral frontal cortex correlated with prediction strength. (A-C) Population time-frequency power during delay between sound and greeble image for (A) HP, (B) MP and (C) NP trials. All plots aligned to greeble onset. (D-F) Population comodulograms for (D) HP, (E) MP, and (F) NP trials during delay. Inset at top left shows dorsolateral frontal electrodes in an example participant. (G-L) Probability distribution of alpha traveling wave directions across delay for two exemplar participants. (G, J) HP, (H, K) MP and (I, L) NP trials. Inset at top right shows task structure with blue square highlighting time window of analysis. A, sound cue; V, greeble image.

**Figure S2.**
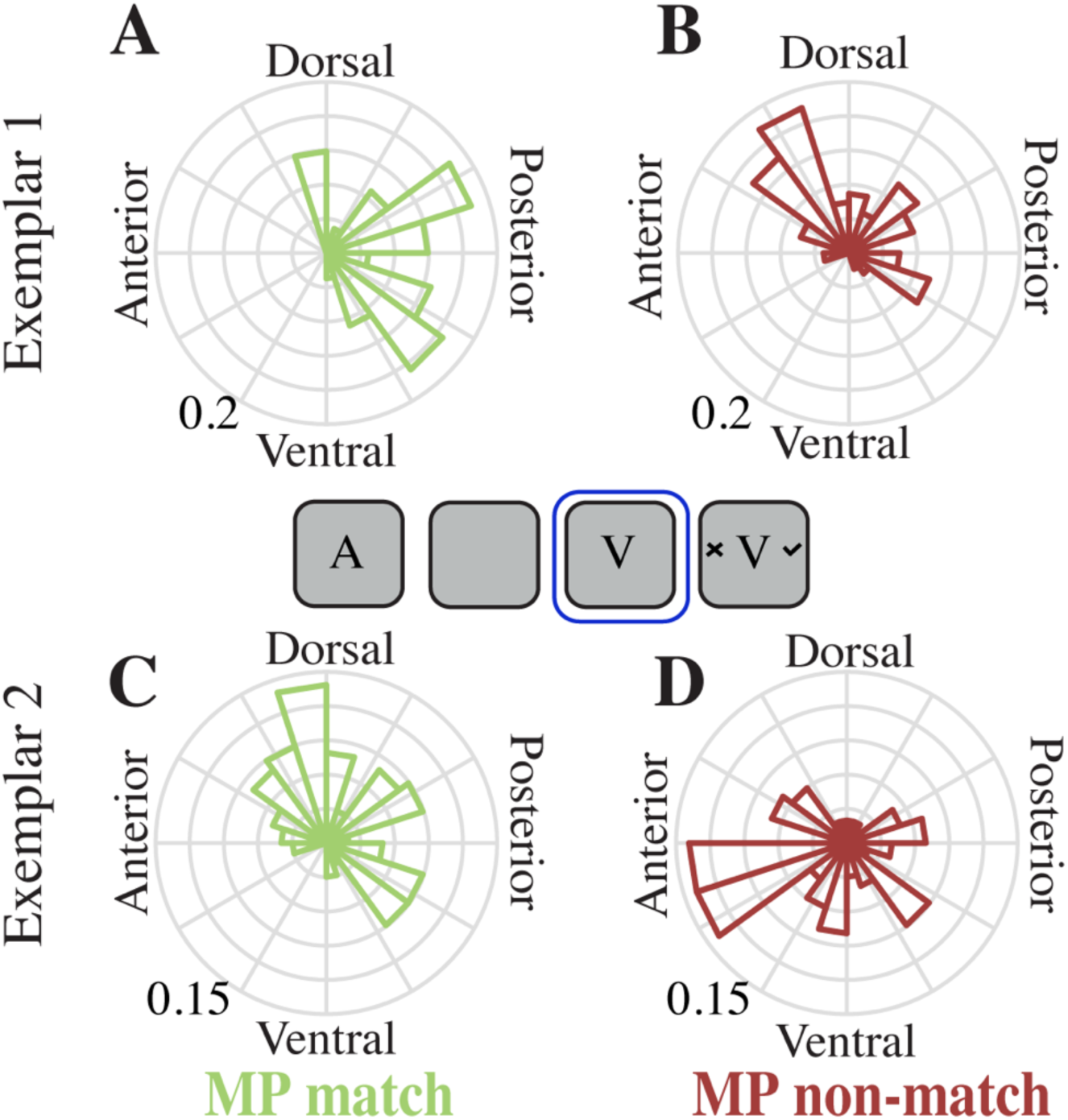
Posterior-to-anterior alpha traveling wave examples after prediction error. Probability distribution of alpha traveling wave directions across period between image and response options for two exemplar participants. (A, C) MP match and (B, D) MP non-match trials. Inset shows task structure with blue square highlighting time window of analysis. A, sound cue; V, greeble image.

**Figure S3.**
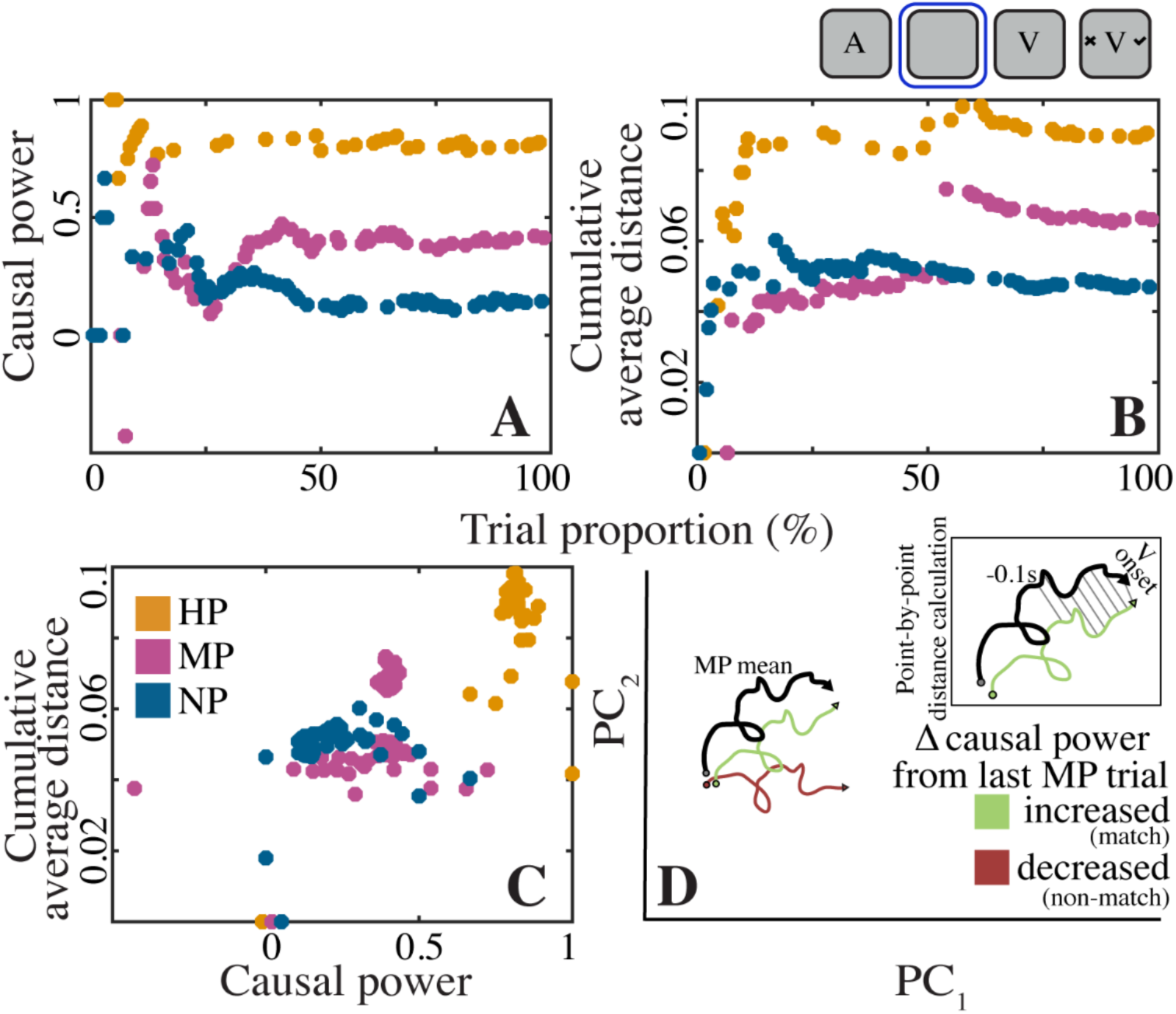
Cumulative average distance correlated to causal power on a trial-by-trial basis. (A) Causal power, and (B) cumulative average distance from the initial trial for each condition (HP, MP, NP) as a function of trial proportion, illustrated for an exemplar participant. Inset at top right shows task structure with blue square highlighting time window of analysis. A, sound cue; V, greeble image. (C) Correlation plot illustrating the relationship between causal power and cumulative average distance for the exemplar participant shown in A-B. We present this example participant to highlight a scenario where, in early trials, A2’s (MP’s) causal power was lower than that of A3 (NP) due to a pseudorandomly generated stimulus presentation sequence (accordingly, the sequence differed for each participant). The neural trajectories represented in (B) closely mirrored causal power changes in (A), with cumulative average distance for early A2 (MP) trials being lower than that of A3 (NP) trials, emphasizing a direct correspondence between causal power and the trajectories. (D) Schematic showing neural trajectories of MP trials influenced by changes in causal power from the preceding MP trial. Here, decreased causal power (red curve) indicates a non-match between the prediction and sensory stimulus in the preceding trial; increased causal power (green curve) indicates a match in the preceding trial. Circle marks starting point, and arrow indicates final position (at greeble onset), of the trajectory. Inset illustrates approach to computing the point-by-point distance of the “MP mean” trajectory (black curve) from an individual MP trial’s trajectory, for the 0.1s before greeble onset.

**Table S1.** Individual participant information.

| **Subject id** | **Age** | **Sex** | **Seizure locus** | **Number of electrodes**<br>**(frontal,**<br>**association**<br>**auditory,**<br>**lateral**<br>**temporal)** |
| --- | --- | --- | --- | --- |
| 1 | 30 | F | Left medial<br>temporal | 96 |
| 2 | 22 | F | Left medial<br>temporal | 21 |
| 3 | 45 | F | Left medial<br>temporal | 37 |
| 4 | 33 | M | Left transverse<br>temporal | 46 |
| 5 | 51 | F | Left medial<br>temporal | 24 |
| 6 | 54 | M | Left medial<br>temporal | 52 |

